# Cell-type-specific transposable element transcription tracks symbiosis and calcification programs in the reef-building coral *Acropora hemprichii*

**DOI:** 10.64898/2026.05.21.726757

**Authors:** Huawen Zhong, Migle Kotryna Konciute, Juntong Hu, Jessica Menzies, Guoxin Cui, Manuel Aranda

**Author notes:** Corresponding author: Guoxin Cui Manuel Aranda.

## Abstract

Transposable elements (TEs) are pervasive components of eukaryotic genomes and major drivers of genome evolution, yet their contribution to cell-type-specific regulatory landscapes remains poorly understood, particularly in non-model marine invertebrates. Here, we integrated single-cell RNA sequencing with pseudo-aligned TE expression profiling to examine how TE transcription relates to cell type identity in the reef-building coral *Acropora hemprichii*. We constructed a cell atlas comprising 4,716 cells across eight major cell types. Notably, TE expression alone was sufficient to accurately resolve all major cell types, indicating that cell-type-specific transcriptional states are robustly reflected in TE activity patterns. We identified 9,759 expressed TEs, of which 333 exhibited strong cell-type-specific activity. These differentially expressed TE features were associated with nearby expressed genes and transcription factor loci, suggesting a relationship between cell-type-specific TE activity and local gene regulatory programs. Genes associated with cell-type-specific TEs were enriched for core coral physiological processes, including calcification, metabolite transport, and symbiosis-related functions. Together, these findings indicate that TE transcription is structured along coral cell-type identity and physiological specialization. Our study provides a single-cell-resolved framework for investigating TE-gene relationships in early-diverging metazoans and a community resource for future functional interrogation in reef-building corals.

## Introduction

Transposable elements (TEs) are DNA sequences that can move within and between genomes, playing a major role in genome evolution and regulatory innovation^1, 2^. Beyond their capacity to reshape genome architecture, TEs can introduce alternative promoters, enhancers, and transcription factor binding sites^3–7^, thereby influencing gene expression programs across development, physiology, and disease^8–12^. Accumulating evidence from model systems indicates that TEs can be co-opted into gene regulatory networks, yet the extent to which TE activity is organized in a cell-type-specific manner remains poorly understood.

A central limitation in studying TE regulation has been the reliance on bulk transcriptomic approaches, which obscure cell-type-specific transcriptional states. As a result, it remains unclear whether TE expression simply reflects background transcriptional noise or whether it encodes structured regulatory information linked to cellular identity and function. Addressing this question requires analytical frameworks capable of resolving TE activity at single-cell resolution and integrating it with gene regulatory programs.

Single-cell RNA sequencing (scRNA-seq) provides a powerful opportunity to overcome this limitation by capturing transcriptional heterogeneity across individual cells^13–15^. While scRNA-seq has been widely applied to define cell types^16–18^ and gene expression programs^19–21^, TE-derived transcripts are often ignored or filtered during standard analyses. Recent advances in pseudo-alignment and TE-aware quantification now enable systematic interrogation of TE transcription from single-cell datasets, opening new avenues to investigate the regulatory landscape of TEs in complex tissues.

Recent work in mouse and human has shown that TE-aware quantification can resolve cell-type structure from scRNA-seq data^22, 23^, indicating that TE transcription carries cell-type-informative signal at single-cell resolution. Whether this holds in invertebrates, and in lineages whose defining cell biology has no vertebrate counterpart, such as cnidarian symbiosis and calcification, remains untested.

Reef-building corals represent an informative system in which to explore cell-type-specific TE activity. As early diverging metazoans with complex cell-type organization and intimate symbioses, corals provide a unique evolutionary context for understanding how mobile genetic elements interface with gene regulatory networks. However, the contribution of TEs to coral cellular identity and functional specialization remains largely unexplored.

In this study, we integrated scRNA-seq with TE expression profiling to investigate the relationship between TEs and cell type identity in the reef-building coral *Acropora hemprichii*. By constructing a single-cell atlas encompassing major cell types and systematically mapping cell-type-specific TE activity, we reveal that TE expression patterns are highly structured and closely associated with gene regulatory programs. Our analyses indicate that differentially expressed TEs are frequently embedded within or proximal to cell-type marker genes and transcription factor loci, indicating that TE transcription is structured along coral cell-type identity and physiological specialization.

## Results

### A high-quality *Acropora hemprichii* reference genome

To facilitate cell-type-resolved gene and TE analyses, we generated a contiguous *de novo* reference genome for *A. hemprichii* from sperm DNA collected during induced spawning in the KAUST *ex situ* Coral Spawning, providing a haploid, symbiont-free source. PacBio HiFi sequencing generated 62.6 Gb of high accuracy reads (3.4M reads; N50 = 19.3 kb; mean Q = 27.6), corresponding to ∼155× coverage of the estimated haploid genome. *De novo* assembly with hifiasm yielded a primary assembly of 489.7 Mb across 176 contigs, with an N50 of 13.8 Mb and a maximum contig length of 26.4 Mb (**Supplementary Table 1**). BUSCO analysis against the metazoa_odb10 lineage recovered 93.2% of complete orthologs, of which only 1.5% were duplicated, consistent with the haploid origin of the template (**Supplementary Fig. 1**). Gene annotation yielded 36,154 protein-coding genes. RepeatModeler^24^ identified 54,265 TE consensus sequences. Genome-wide RepeatMasker^25^ annotation showed that repetitive elements covered 66.56% of the genome, with DNA transposons, LINEs, LTRs, and unclassified repeats representing the dominant TE classes (**Supplementary Fig. 2**). This assembly and its annotations were used in all downstream single-cell gene and TE expression analyses.

### A single-cell atlas defines major cell types in *Acropora hemprichii*

To establish a cellular framework for investigating TE activity, we generated a single-cell transcriptomic atlas of *A. hemprichii* from three individuals (**Fig. 1a**). After quality filtering and integration, we obtained a total of 4,716 cells with 33,509 detected genes (**Supplementary Table 2**). Unsupervised clustering resolved ten transcriptionally distinct cell clusters.

**Fig. 1.**
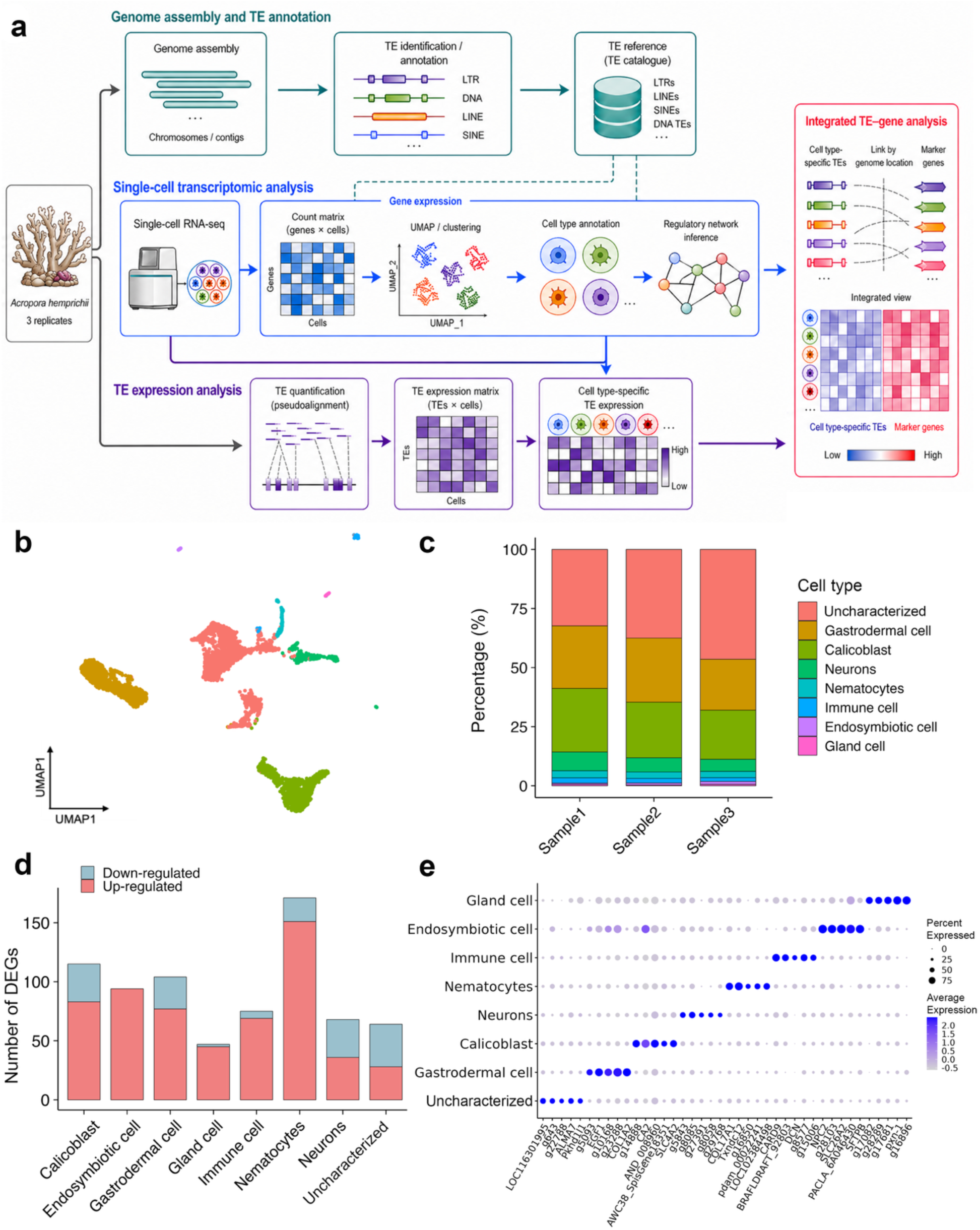
*Acropora hemprichii cell type atlas.* **a.** Overview of the project; **b.** UMAP plot for the identified eight cell types; **c.** Cell type proportion for three samples; **d.** Bar plot of the number of differentially expressed genes for eight cell types; **e.** Dot plot for top five marker genes in eight cell types.

To annotate these clusters, we first examined symbiont-derived UMI counts per cell by mapping scRNA-seq data to the *Cladocopium* genome^26^, the predominant symbiont genus in *A. hemprichii*. One cluster showed markedly higher algal UMI counts and was annotated as the endosymbiotic cell type (**Supplementary Fig. 3**). The remaining clusters were then annotated by cross-species mapping with SAMap^27^ to a published single-cell atlas of the stony coral *Stylophora pistillata*^28^ (**Supplementary Fig. 4**). SAMap mapping confirmed the endosymbiotic assignment and resolved six additional clusters to individual *S. pistillata* cell types, including gastrodermal cells, calicoblasts, neurons, nematocytes, immune cells, and gland cells (**Supplementary Fig. 4**). The remaining three clusters showed diffuse mapping across multiple *S. pistillata* cell groups, indicating a lack of clear one-to-one correspondence, and were merged into a single uncharacterized group. Together, these assignments defined seven annotated cell types plus one uncharacterized population (**Fig. 1b**), recovered across all three individuals with comparable proportional representation (**Fig. 1c**).

Marker gene analysis identified 752 cell-type-specific genes across the atlas (**Fig. 1d**). Canonical cnidarian cell markers were recovered in their expected clusters, including bicarbonate transporter SLC4A4 and carbonic anhydrase CA2 in calicoblasts, and cholesterol transporter NPC2 and ammonium transporter Rh in endosymbiotic cells (**Fig. 1e, Supplementary Table 3**).

To identify the transcriptional regulators underlying these cell identities, we inferred gene regulatory networks with SCENIC^29^. The top five regulons per cell type revealed distinct sets of driver transcription factors (TF; **Fig. 2a**). Consistent with this high specificity, TFs of the same cell type interacted more intensively among themselves than across cell types (**Fig. 2b**). On average, within-cell-type regulon connectivity was ∼2× stronger than cross-cell-type connectivity, indicating that each cell type is underpinned by a largely self-contained regulatory program.

**Fig. 2.**
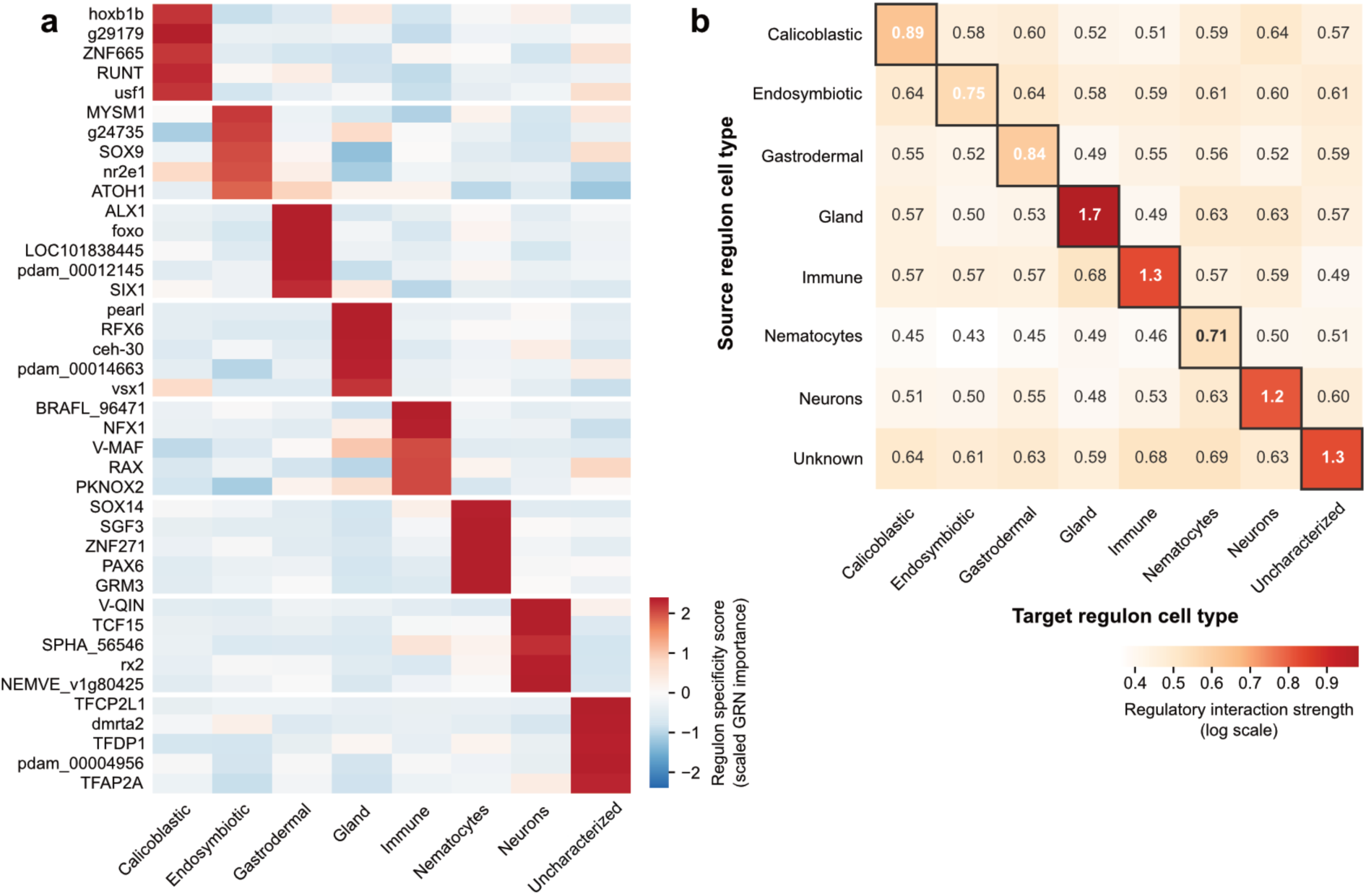
*Cell type specific regulons in Acropora hemprichii.* **a.** Heatmap plot of regulon specificity score for the top five regulons per cell type, scaled across cell types. Color scale indicates scaled GRN importance. **b.** Pairwise matrix of mean regulatory interaction strength between cell types, computed from the full SCENIC-inferred gene regulatory network. Each entry represents the mean edge importance from TFs assigned to the source cell type (rows) to target genes assigned to the target cell type (columns). Diagonal entries (outlined) correspond to within-cell-type regulation. Values shown are the mean edge importance per cell-type pair. Color scale log-transformed for visualization.

### Single-cell TE expression is cell-type-specific

To examine TE activity at single-cell resolution, we quantified TE expression from our scRNA-seq data using a pseudo-alignment strategy optimized for repetitive elements^30^. After filtering out TEs detected in fewer than three cells, 9,759 expressed TEs were retained for downstream analyses. Clustering based solely on the expression of the 2,000 most variable TEs recovered the same eight cell populations resolved by gene expression (**Fig. 3a**), indicating that TE transcription captures structured, cell-type-specific information rather than stochastic background noise. This pattern was consistent across cells with varying total transcript counts, arguing against a primary effect of library size or global transcriptional activity (**Supplementary Fig. 5, Supplementary Table 4**). Across all cell types, the median number of detected TEs per cell scaled linearly with the number of detected genes (**Fig. 3b**), suggesting that TE transcription is broadly integrated with cellular transcription output. Differential expression analysis identified 333 TEs with cell-type-specific expression patterns (DETEs; **Fig. 3c**). Uncharacterized TEs, putative coral-specific elements detected as repeats but lacking homology to annotated TE families, formed the largest DETE class in every cell type. Among classified families, DNA transposons and long interspersed nuclear element (LINE) contributed most of the remaining DETEs, while long terminal repeat (LTR) and short interspersed nuclear element (SINE) retrotransposons were largely absent across cell types.

**Fig. 3.**
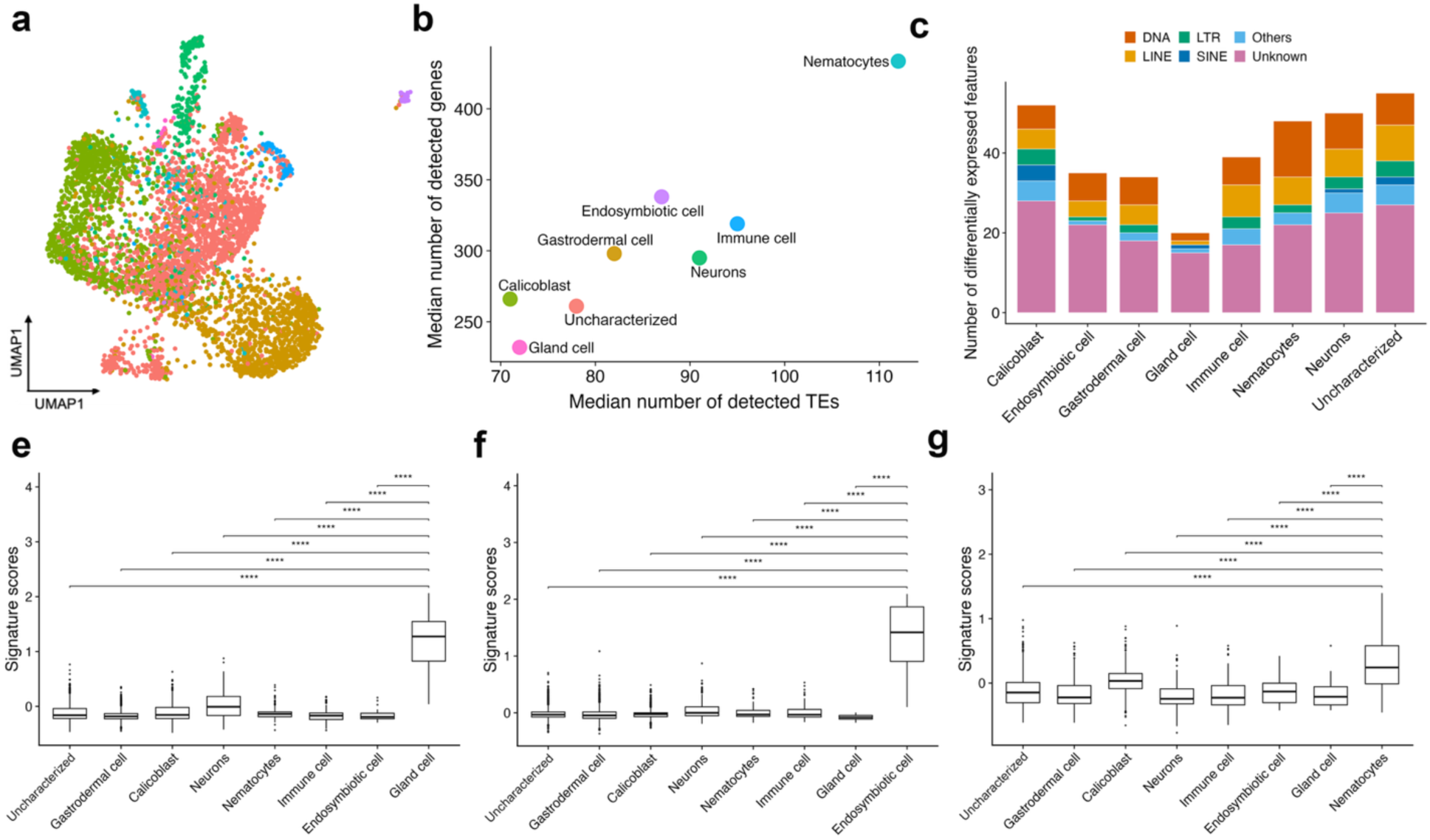
*Single-cell TE expression distinguishes cell populations in Acropora hemprichii.* **a.** UMAP plot based on 2000 most variable individual TE expressions. Cells are color-coded based on the cell population; **b.** Median number of detected TE and genes for each cell type; **c.** Bar plot of a total of 333 DETEs’ family category for each cell type. DNA: DNA transposons, LINE: long interspersed nuclear element, LTR: long terminal repeats, SINE: short interspersed nuclear elements, Others: rolling-circles, small RNA, satellites, and simple repeats; **d-f**. The signature scores for gland cell, endosymbiotic cell and nematocytes, respectively. A pair-wise t-test comparison is conducted between the target cell types and the others. ***: p-value <0.001, **: p-value < 0.01, *: p-value <0.05.

To further assess the specificity of DETE expression, we next calculated DETE signature scores for each cell type (**Fig. 3e-g, Supplementary Fig. 6**). DETEs from representative cell types, including gland cell (**Fig. 3d**), endosymbiotic cell (**Fig. 3e**), and nematocytes (**Fig. 3f**), clearly discriminated their corresponding populations from the rest of the atlas. Together, these results show that individual TE expression profiles are sufficient to delineate coral cell types at the single-cell resolution.

### Cell-type-specific TE expression is closely associated with nearby gene activity

The concordance between TE- and gene-based cell-type resolution suggested a close relationship between TE transcription and gene expression. To further examine this association, we restricted the analysis to positive DETEs and DEGs with adjusted *P* value less than 0.05, yielding 261 DETEs and 1,872 DEGs. We then calculated Pearson correlation coefficients using cell-type-averaged expression profiles. DETEs and DEGs of the same cell type showed consistent positive correlated across the atlas (**Fig. 4a**), with endosymbiotic and gland cells showing the strongest coefficients.

**Fig. 4.**
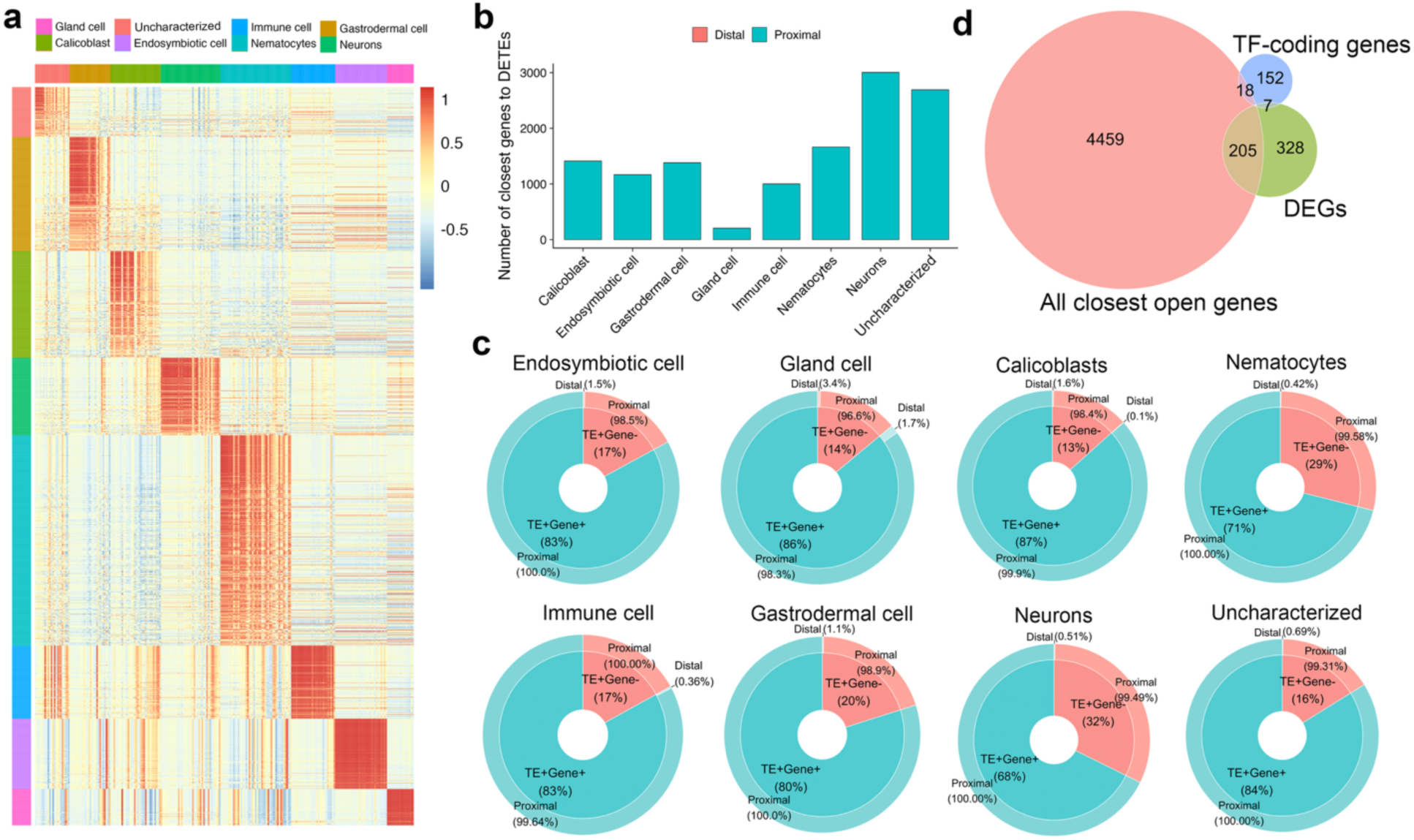
*DETEs reveal the cell type signatures.* **a.** Heatmap of the identified 261 DETEs and 1872 DEGs expression based on scaled data. Positive DEGs and DETEs are selected with p-adjusted-value < 0.05. **b.** Bar plot for the number of closest genes amount to the DETE for each cell type; **c.** Donut plots show if the closest genes are open or not and their distance to the DETEs. Distal: distance between the DETE and its closest genes > 2000 bp. Proximal: distance between the DETE and its closest genes < 2000 bp. **d.** Venn diagram showing the intersection among all the closest open genes, all unique DEGs and TF-coding genes.

The co-variation between DETE and DEG expression raised the question of whether genomic proximity contributes to this relationship. Because TE expression was quantified at the TE family level and each DETE family is represented by multiple genomic copies, we examined DETE–gene proximity at several levels of organization. At the TE family level, 93.5% of DETEs possessed at least one copy within 2 kb of a gene, compared with 72.3% of non-DETE families (Fisher’s exact test OR = 5.49, p = 7.25 × 10⁻¹⁶), indicating that cell-type-specific TE families are significantly enriched for gene-proximal copies. Similar but more modest enrichments were observed at the individual locus and TE family–gene pair levels (**Supplementary Fig. 7**). Together, these results suggest that DETE transcription is non-randomly associated with gene-proximal genomic regions.

To further relate this proximity pattern to cell-type-specific TE activity, we next assigned each cell type–specific DETE to its nearest gene and retained the shortest-distance gene for each cluster–TE pair. This yielded 12,537 DETE-associated closest-gene assignments, of which 12,510 were located within 2 kb of the corresponding DETE locus (99.78%; **Fig. 4b**). In contrast, non-DETE-associated cluster–TE assignments showed a significantly lower proportion of proximal gene associations, resulting in a strong enrichment for gene-proximal associations among DETEs (Fisher’s exact test, OR = 30.69, p = 1.16 × 10⁻²⁹⁷. **Supplementary Fig. 8**). These results further support that cell-type-specific TE transcription is preferentially associated with nearby genes, consistent with a potential contribution of local genomic context to TE–gene co-variation.

To ask whether this proximity reflects a functional link, we checked whether each DETE’s nearest gene was itself expressed in the matching cell type. This was the case for 78% of DETEs, while the remaining 22% had nearest genes that fell below the detection threshold (**Fig. 4c**), indicating that cell-type-specific TE transcription is consistently associated with local gene activity.

### DETEs preferentially localize to transcriptional regulators underlying cell-type identity

We further examined the expressed closest genes (“open closest genes”) to determine their relationship with cell-type marker genes and transcription factors (**Fig. 4d**). Of these open closest genes, 38% were cell type markers of the same cell type as their associated DETEs. We then categorized the open closest genes into four groups: (1) DEGs specific to the same cell type, (2) TF-coding genes, (3) genes that were both TF-coding and differentially expressed, and (4) TF-coding genes whose regulon targeted cell-type-specific DEGs.

Across all cell types, we identified 88 unique DEGs that were closest to DETEs within the same cell type. Since individual DEGs could be associated with multiple DETEs, we quantified DETE-DEG pairings and their genomic configurations (**Fig. 5b, c**). Remarkably, the majority of DETEs were embedded within gene bodies of their associated DEGs, consistent with a tight genomic coupling between TE insertions and host gene transcription.

**Fig. 5.**
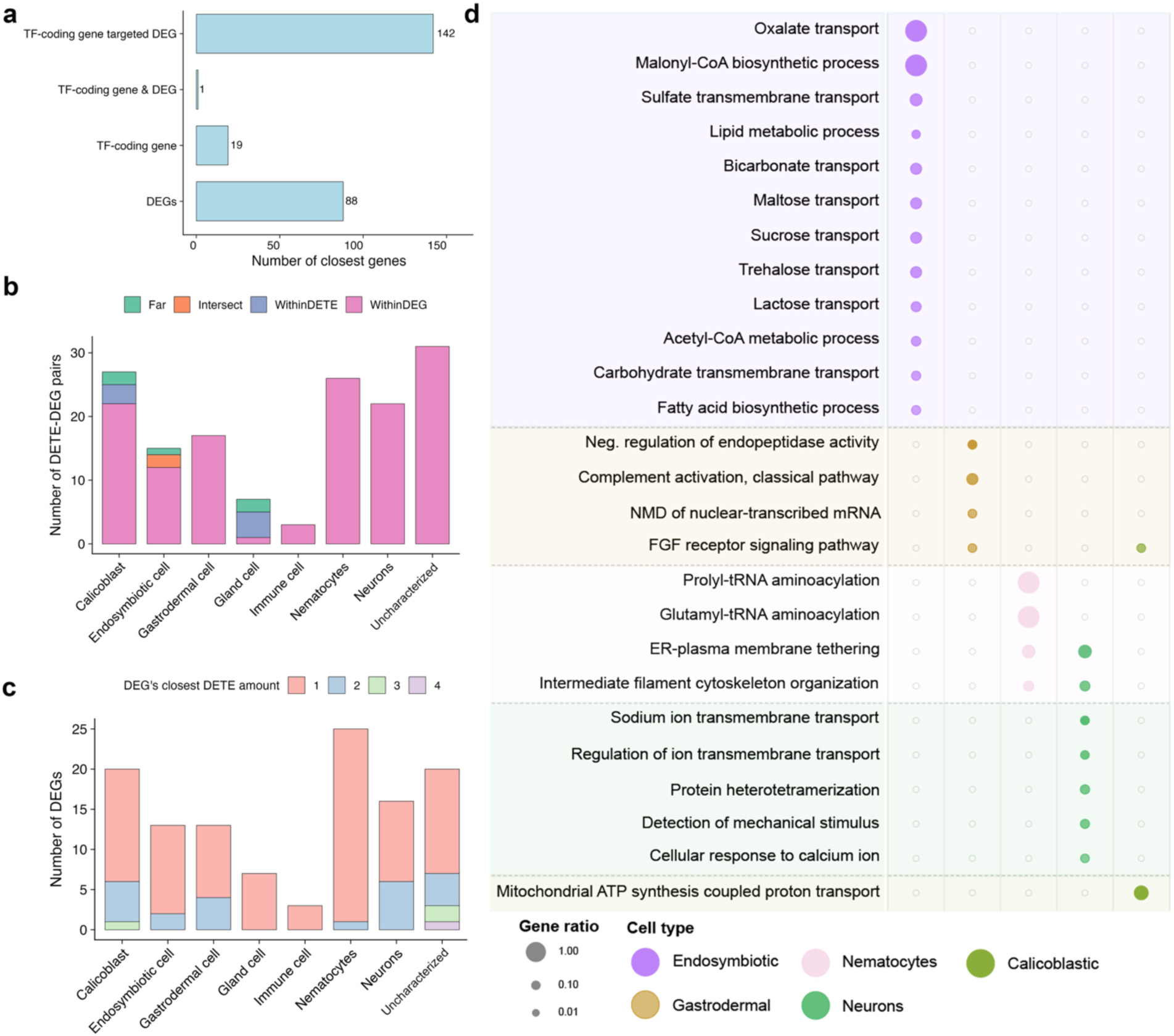
*Closest gene to the DETEs categories.* **a.** Bar plot for the number of closest genes to DETEs based on different filtering criteria. DEGs represent the closest genes of DETEs are the DEGs for the same cell type; TF-coding genes represent the closest genes are TF-coding genes; TF-coding gene & DEGs represent the closest gene is TF-coding genes and DEGs at the same time; TF-coding gene targeted DEG represents the closest gene is TF-coding gene whose target gene is the DEG for the same cell type. **b.** Bar plot of DETE-DEGs pairs in 8 cell types with their distance categories. **c.** Bar plot for the unique DEGs in 8 cell types with their closest DETE number. **d.** Enriched biological processes in the main cell types of *A. hemprichii*.

Functional enrichment analysis of DETE-associated DEGs revealed strong enrichment for biological processes characteristic of specific cell types, particularly in endosymbiotic, calicoblastic, and gastrodermal cells (**Fig. 5d**). For example, the enriched biological processes in endosymbiotic cells exhibited respiratory gaseous exchange, bicarbonate transport, and sugar transport, suggesting that the DETE-harboring genes might play essential roles in cellular function.

Interestingly, all TF-coding genes identified as closest to DETEs contained TEs within their gene bodies. In total, 37 such TF-coding genes targeted downstream DEGs in a cell-type-specific manner. This observation suggests that TE insertions may influence cellular identity not only through direct association with marker genes, but also indirectly by modulating transcriptional regulons that coordinate broader gene expression programs.

## Discussion

This study provides the first single-cell-resolved view of transposable element (TE) activity in corals and demonstrates that TE transcription is structured in a cell-type-specific manner rather than reflecting passive or stochastic genomic noise. TE expression alone was sufficient to distinguish all major cell populations, extending to a non-vertebrate, early-diverging metazoan an observation previously made in mouse and human^22, 23^ and positioning TEs as informative features of cellular identity in cnidarians.

A central finding is the close genomic and transcriptomic association between DETEs and cell-type-specific gene programs. Nearly all DETEs were located proximal to marker genes, with many embedded within their gene bodies. This genomic arrangement is consistent with models in which TE insertions can function as cis-regulatory elements that modulate transcriptional output of neighboring genes in a cell-type-specific manner. In addition to these local associations, DETEs were frequently embedded in TF-coding loci whose regulons define cell-type programs, suggesting an indirect route by which TE insertions may influence broader gene regulatory networks. Such multi-level positioning mirrors observations in vertebrates, where TE-derived sequences contribute regulatory motifs, reshape TF binding landscapes, and promote regulatory innovation^4, 31, 32^. We note that the associations between TE expression, genomic proximity, and gene regulatory programs are correlative and do not imply direct cis-regulatory activity, which will require targeted functional validation.

The ability of TE-based expression profiles to recapitulate gene-defined cell types underscores that TE activity represents an informative dimension of the single-cell transcriptome. Standard scRNA-seq pipelines often down-weight or discard repetitive reads, which can obscure biologically meaningful variation. Our results indicate that, at least in the TE-rich coral genome, these reads encode robust cell identity information. Incorporating TE expression into single-cell analyses therefore has the potential to sharpen cell type classification, reveal cryptic cellular states, and expose regulatory mechanisms that would remain hidden when focusing solely on protein-coding genes.

Functional enrichment of DETE-associated genes further supports a biologically meaningful role for TEs in coral cell biology. DETE-proximal genes were enriched for processes central to coral physiology, including calcification, gas exchange, and metabolite transport. In endosymbiotic cells, DETE-linked genes were particularly enriched in bicarbonate transport and sugar exchange pathways, which are directly relevant to metabolic interactions between coral hosts and their dinoflagellate symbionts. These patterns are consistent with co-option of TE-derived sequences into regulatory circuits supporting symbiosis-related functions and align with growing evidence that TEs facilitate evolutionary innovation by rewiring regulatory networks under ecological and environmental pressures^33–38^.

Family-level biases in DETE composition suggest that not all TE lineages contribute equally to regulatory architecture. Most DETEs originated from DNA transposons and LINEs, with some retrotransposon classes, e.g. LTRs or SINEs, being underrepresented or absent among DETEs in several highly specialized cell types. Such non-random family-level patterns may reflect differences in insertion history, chromatin accessibility, epigenetic regulation, or intrinsic capacities of TE lineages to donate promoters, enhancers, or splice sites. Dissecting these contributions will require finer-scale annotation of TE subfamilies, their evolutionary ages, and the associated epigenomic contexts within coral genomes.

Integration of TE expression with inferred gene regulatory networks (GRNs) provides additional insight into how TEs may influence cell-type specification. DETEs were frequently intersected TF-coding genes whose regulons contain cell-type marker genes, suggesting that TE insertions may act at multiple hierarchical levels: locally tuning expression of effector genes while simultaneously modulating central regulatory nodes. In an evolutionary context, such architectures could enable incremental diversification of cell states, for example, refining endosymbiotic or calcifying cell states, while preserving conserved ancestral regulatory frameworks shared across cnidarians.

The findings also highlight TE activity as a potential source of regulatory plasticity in the face of environmental change. Reef-building corals are exposed to escalating stressors, including ocean warming and acidification, that challenge physiological homeostasis. TE-derived regulatory modules, such as novel enhancers, alternative promoters, or modifications to transcription factor loci, could facilitate rapid diversification of cellular responses involved in oxidative stress responses, metabolic flexibility, and biomineralization. Observations in other taxa linking TE activation to stress responses and developmental transitions suggest that similar mechanisms may contribute coral resilience by reshaping regulatory landscapes at the single-cell level ^33–38^.

Several limitations should be considered. Quantification of TE expression from scRNA-seq is intrinsically challenging due to multi-mapping reads, high sequence similarity among young TE copies, and limited sequencing depth. The pseudo-alignment strategy employed here offers a practical solution for capturing TE-level expression patterns but does not fully resolve locus-specific contributions, particularly for recent TE insertions. The diversity and dynamics of active TEs are therefore likely to be underestimated. A further consideration is that part of the DETE signal could reflect co-transcription with neighboring host genes rather than independent TE expression, particularly for DETEs embedded within gene bodies. Family-level pseudo-alignment cannot fully disambiguate these scenarios; locus-resolved, strand-specific quantification, ideally combined with chromatin accessibility or nascent transcription data, will be required to definitively partition independent TE activity from host-gene read-through. The observation that TE-only clustering recapitulates cell-type structure across cells with comparable total transcript counts (Supplementary Fig. S5) argues against a uniform read-through artefact, but does not rule out a partial contribution at the level of individual DETE-gene pairs. Moreover, the relationships between DETEs, marker genes, and transcriptional regulators are correlative. Establishing causality will require targeted functional perturbations, such as CRISPR-mediated deletion or epigenetic editing of specific TE insertions, as well as reporter assays testing TE-derived regulatory sequences in coral cells or tractable heterologous systems. Besides, comparative analyses across additional cnidarian species and integration of epigenomic data layers, including chromatin accessibility, DNA methylation, and histone modification, will further clarify the generality and mechanistic basis of TE-associated regulation.

In summary, this study identifies TEs as an informative and previously underappreciated dimension of the coral single-cell transcriptome, with cell-type-specific TE expression patterns positionally and transcriptionally associated with gene programs underlying symbiosis, calcification, and physiological specialization. More broadly, explicit consideration of TE biology in single-cell genomics may refine our understanding of gene regulation and evolutionary innovation across diverse metazoan lineages, particularly in organisms with TE-rich genomes and dynamic ecological challenges.

## Methods

### Coral spawning, sperm collection, and genome sequencing

Parent colonies of *A. hemprichii* were collected in November 2021 from central Red Sea reefs near KAUST (22°18′13.8″N 38°57′51.0″E, 5–15 m depth) and maintained in the KAUST *ex situ* Coral Spawning System^39^ with programmed temperature, photoperiod, and lunar cycles. The day–night cycle was inverted six weeks before anticipated spawning to enable daytime gamete collection. Spawning colonies were transferred to isolated buckets at the first signs of setting, where released gamete bundles were collected using transfer pipettes. Bundles were dissociated by gentle transfer between beakers, sperm was siphoned away from eggs and retained for DNA extraction. High-molecular-weight DNA was extracted with the Monarch Genomic DNA Purification Kit for Cells & Blood (New England Biolabs), quantified with the Qubit dsDNA Broad Range assay (Thermo Fisher Scientific), and assessed on the Femto Pulse platform (Agilent). Sequencing was performed on the PacBio Sequel II system with HiFi library preparation. HiFi reads were assembled with hifiasm^40^ v0.18 under default parameters. Assembly completeness was assessed with BUSCO^41^ v5.1.2 against the metazoa_odb10 lineage. The resulting primary assembly was used for all downstream analyses.

### Gene model prediction and functional annotation

Prior to gene prediction, the assembly was soft-masked in two rounds. Known metazoan repeats were first masked with RepeatMasker^25^ (v4.1.0) using the metazoa repeat library. Species-specific repeat families were then identified *de novo* with RepeatModeler^24^ (v2.0.1) including LTR structural discovery, and the resulting library was used in a second RepeatMasker round to soft-mask the previously masked genome. Gene models were predicted with BRAKER2^42^ (v2.1.5) using the soft-masked assembly.

Predicted protein sequences were functionally annotated using a customized pipeline (https://github.com/lyijin/annotating_proteomes), as previously applied to several cnidarian genomes^43, 44^. For each gene model, the longest predicted protein isoform was queried against three databases by BLASTP (NCBI BLAST+ v2.15.0; e-value cutoff 1e-5): UniProt/Swiss-Prot, UniProt/TrEMBL, and NCBI nr. The top 20 hits were retained for Swiss-Prot and TrEMBL, and the top hit for nr. For Swiss-Prot and TrEMBL hits, the highest-ranked match with associated Gene Ontology (GO) annotations was selected as the representative hit, using the GO annotation database (goa_uniprot_all.gaf) for cross-referencing. Each gene model was assigned a textual annotation following the priority Swiss-Prot > TrEMBL > nr, and GO terms were assigned via the go-basic.obo hierarchy. The resulting annotation was used as the reference for downstream analyses.

### Single-cell RNA sequencing

*A. hemprichii* (n = 3) fragments were collected by dissecting the adult colonies with a cutter into fragments sized ∼2 cm. Fragments were incubated in 10 ml of 0.22 μm-filtered Ca/Mg-free artificial seawater (23 g/L NaCl; 0.763 g/L KCl; 3 g/L Na_2_SO_4_; 0.25 g/L NaHCO_3_; pH 8.0) for 2 hours on a Belly Dancer at 30 rpm. Coral cells were then dissociated by gently scraping and peeling off the tissue using a 10 μl sterile pipette tip. The cell suspension was then filtered twice through a 40 μm cell strainer (Thermo Fisher Scientific). The filtered cell suspensions were stained with 2 μl Calcein AM working solution and sorted for live cells using FACS before being loaded directly on the 10X Chromium system (10X Genomics). Single-cell libraries were generated using the 10X Chromium Single-Cell 3′ Reagent Kit V3 according to the manufacturer’s instructions. Library size distribution and concentration were determined using the Agilent Bioanalyzer with the High Sensitivity DNA kit and the QuantStudio 3 real-time polymerase chain reaction system (Thermo Fisher Scientific) with the KAPA DNA quantification kit (Roche), respectively. The libraries were sequenced using an Illumina HiSeq 4000 to generate paired end reads.

### scRNA-seq analysis

#### Data processing

We used CellRanger^45^ (v.5.0.1) to generate raw gene expression matrices for each sample. Next, we used R software (v.4.0.4) with the Seurat package^46^ (v.4.0.3) to analyze the output filtered gene expression matrices. Cells were filtered to include those with unique molecular identifiers (UMI) counts ≥200 and < 99% of the highest UMI count in the dataset, thereby excluding low-coverage cells and extreme high-UMI outliers. Then we used *NormalizeData* to normalize the data, followed by calculating the 2000 features with high cell-to-cell variation through the *FindVariableFeatures* function. We further used *FindIntegrationAnchors* function and *IntegrateData* function to integrate all the samples into one dataset followed by using the *Scale* function to scale the integrated dataset. Next, we used *RunPCA* and *RunUMAP* functions to reduce the dimensionality of the dataset. Finally, we used *FindNeighbors* and *FindClusters* functions to cluster cells.

#### Cell type annotation

SAMap^27^ is a tool that aligns cell atlases in two mutually reinforcing directions, mapping both the genes and the cells, with each feeding back into the other. It accounts for the complexity of gene evolution and is able to map single-cell atlases between evolutionarily distant species. We followed SAMap’s tutorial (https://github.com/atarashansky/SAMap/blob/main/SAMap_vignette.ipynb) to run the pipeline. Specifically, we mapped between the clusters in *Acropora hemprichii* and the cell types of *Stylophora pistillata*^28^. Alignment threshold was set as 0.2 as default.

#### DEG identification and functional annotation

To identify cluster-specific differentially expressed genes, we used Seurat *FindAllMarkers* with *only.pos = FALSE, min.pct = 0.25* and *logfc.threshold = 0.25*. The marker genes are further filtered with adjusted p-value < 0.05. The signature score for the markers is calculated using the Seurat function *AddModuleScore*.

Differentially expressed genes used in the downstream correlation analysis with the differentially expressed TE were identified using *FindAllMarkers* with *only.pos = TRUE*. We filtered the DEGs whose adjusted p-value < 0.05. The same settings were performed for the differentially expressed TE. GO term enrichment analysis for the DEGs was performed using the R software TopGO package^47^.

#### Gene regulatory network analysis

We used SCENIC^29^ to infer the transcription factors and gene regulatory networks following the tutorial(https://github.com/aertslab/SCENICprotocol/blob/master/notebooks/PBMC10k_SCENIC-protocol-CLI.ipynb). In brief, we generated the orthologous transcription factor *A. hemprichii*, based on the published TF of *Nemastella vectensis*^48^ using orthoFinder^49^. We used the orthologous TF with the single-cell expression data to generate the co-expression matrix and the candidate regulon with the default parameters. We next built up the motif whole-genome ranking database for *A. hemprichii*. To do this, we first generated *A. hemprichii*’s promoter regions fasta file, which selected the 5k bp upstream regions in front of the transcription start site (TSS). We secondly generated the motif list by downloading all the motif data from the Jaspar database^50^. We then constructed the motif2TF database, which links motifs to transcription factors by substituting human gene symbols in the SCENIC-provided human motif2TF file with their homologs from *A. hemprichii*. Next, we built the cisTarget database, calculated the genome ranking, and performed the motif enrichment and TF-regulon prediction following the tutorial. Finally, we used the default parameter to further filter out the genes by the enrichment results.

#### TE identification and expression analysis

We identified the TE sequences through the combination of de novo-based and homology-based methods. We de novo predicted all the TE consensus sequences using LongRepMasker^51^ (v2.1.2) with default settings. We then combined the predicted sequences and the curated homology database from Dfam^52^ , which is subsetted by the Actiniaria family, as the annotation database for *A. hemprichii*. Finally, we ran RepeatMasker^25^ to identify and annotate the TE consensus sequences for *A. hemprichii*.

To obtain the TE expression sparse matrix, we first used the kallisto^53^ tool to perform the pseudoalignment for the raw scRNA sequencing data of *A. hemprichii* for each sample separately. Then, the sparse matrix was generated according to the kallisto tutorial (https://bustools.github.io/BUS_notebooks_R/10xv3.html#sparse_matrix). For each sample, we retained only the cells that were assigned to eight different cell types in the scRNA-seq analysis in Seurat. Then we performed the normalization, integration, dimension reduction, and the cell type marker TEs identification process as described in scRNA-seq data analysis. Finally, we used the bedtools^54^ to find the closest gene to the specific TEs on the same strand.

### Statistical analysis

To assess enrichment of TE–gene proximity, DETE- and non-DETE-associated elements were compared at multiple analytical levels, including genomic locus, TE ID, TE ID–gene ID pair, and cluster–TE closest-gene assignment levels. TE–gene associations were classified as proximal if the TE overlapped a gene or was located within 2 kb of the nearest gene; all other associations were classified as distal. For each level, the proportions of proximal and distal associations were compared between DETEs and non-DETEs using Fisher’s exact test, with odds ratios reported as effect sizes. Differences in TE–gene distance distributions were additionally assessed using the Mann–Whitney U test.

For the cell type–specific DETE proximity analysis, each cluster–TE pair was assigned to its nearest gene by retaining the shortest TE–gene distance. Proximal enrichment among DETE-associated closest-gene assignments was then evaluated relative to non-DETE assignments using Fisher’s exact test. TE signature scores were compared between each target cell type and all other cells using pairwise t-tests. Where applicable, p-values from multiple comparisons were adjusted using the Benjamini–Hochberg method. Statistical significance was denoted as ***p < 0.001, **p < 0.01, and *p < 0.05.

Fisher’s exact tests, Mann–Whitney U tests, and chi-square tests were performed in Python using *scipy.stats*. Pairwise t-tests were performed in R using base *stats* functions.

## Data and code availability

The single-cell data can be downloaded from: [will be provided upon publication]

All the code for analysis is available from https://github.com/huawen-poppy/TE_analysis_of_Acropora.git

## Reference

1. Bourque, G. et al. Ten things you should know about transposable elements. Genome biology 19, 1–12 (2018).

2. Klein, S.J. & O’Neill, R.J. Transposable elements: genome innovation, chromosome diversity, and centromere conflict. Chromosome Research 26, 5–23 (2018).

3. Ayarpadikannan, S. & Kim, H.-S. The impact of transposable elements in genome evolution and genetic instability and their implications in various diseases. Genomics & informatics 12, 98–104 (2014).

4. Chuong, E.B., Elde, N.C. & Feschotte, C. Regulatory activities of transposable elements: from conflicts to benefits. Nature Reviews Genetics 18, 71–86 (2017).

5. Chen, J.-M., Férec, C. & Cooper, D.N. LINE-1 endonuclease-dependent retrotranspositional events causing human genetic disease: mutation detection bias and multiple mechanisms of target gene disruption. Journal of Biomedicine and Biotechnology 2006 (2006).

6. Sobczak, K. & Krzyzosiak, W.J. Structural determinants of BRCA1 translational regulation. Journal of Biological Chemistry 277, 17349–17358 (2002).

7. Chen, L.-L. & Carmichael, G.G. Gene regulation by SINES and inosines: biological consequences of A-to-I editing of Alu element inverted repeats. Cell cycle 7, 3294–3301 (2008).

8. Jang, H.S. et al. Transposable elements drive widespread expression of oncogenes in human cancers. Nature genetics 51, 611–617 (2019).

9. Tam, O.H., Ostrow, L.W. & Gale Hammell, M. Diseases of the nERVous system: retrotransposon activity in neurodegenerative disease. Mobile DNA 10, 1–14 (2019).

10. Göke, J. et al. Dynamic transcription of distinct classes of endogenous retroviral elements marks specific populations of early human embryonic cells. Cell stem cell 16, 135–141 (2015).

11. Grow, E.J. et al. Intrinsic retroviral reactivation in human preimplantation embryos and pluripotent cells. Nature 522, 221–225 (2015).

12. Percharde, M. et al. A LINE1-nucleolin partnership regulates early development and ESC identity. Cell 174, 391–405. e319 (2018).

13. Macaulay, I.C., Ponting, C.P. & Voet, T. Single-cell multiomics: multiple measurements from single cells. Trends in genetics 33, 155–168 (2017).

14. Zhang, Y. et al. Single-cell RNA sequencing in cancer research. Journal of Experimental & Clinical Cancer Research 40, 1–17 (2021).

15. Liang, J., Cai, W. & Sun, Z. Single-cell sequencing technologies: current and future. Journal of Genetics and Genomics 41, 513–528 (2014).

16. Stegle, O., Teichmann, S.A. & Marioni, J.C. Computational and analytical challenges in single-cell transcriptomics. Nature Reviews Genetics 16, 133–145 (2015).

17. Baslan, T. & Hicks, J. Unravelling biology and shifting paradigms in cancer with single-cell sequencing. Nature Reviews Cancer 17, 557–569 (2017).

18. Levitin, H.M., Yuan, J. & Sims, P.A. Single-cell transcriptomic analysis of tumor heterogeneity. Trends in cancer 4, 264–268 (2018).

19. Ren, X., Kang, B. & Zhang, Z. Understanding tumor ecosystems by single-cell sequencing: promises and limitations. Genome biology 19, 1–14 (2018).

20. A single-cell transcriptomic atlas characterizes ageing tissues in the mouse. Nature 583, 590–595 (2020).

21. Chen, G., Ning, B. & Shi, T. Single-cell RNA-seq technologies and related computational data analysis. Frontiers in genetics, 317 (2019).

22. He, J. et al. Identifying transposable element expression dynamics and heterogeneity during development at the single-cell level with a processing pipeline scTE. Nature communications 12, 1456 (2021).

23. Shao, W. & Wang, T. Transcript assembly improves expression quantification of transposable elements in single-cell RNA-seq data. Genome research 31, 88–100 (2021).

24. Flynn, J.M. et al. RepeatModeler2 for automated genomic discovery of transposable element families. Proceedings of the National Academy of Sciences 117, 9451–9457 (2020).

25. Chen, N. Using Repeat Masker to identify repetitive elements in genomic sequences. Current protocols in bioinformatics 5, 4.10. 11–14.10. 14 (2004).

26. Shoguchi, E. et al. Two divergent Symbiodinium genomes reveal conservation of a gene cluster for sunscreen biosynthesis and recently lost genes. BMC genomics 19, 1–11 (2018).

27. Tarashansky, A.J. et al. Mapping single-cell atlases throughout Metazoa unravels cell type evolution. Elife 10 (2021).

28. Levy, S. et al. A stony coral cell atlas illuminates the molecular and cellular basis of coral symbiosis, calcification, and immunity. Cell 184, 2973–2987. e2918 (2021).

29. Van de Sande, B. et al. A scalable SCENIC workflow for single-cell gene regulatory network analysis. Nature Protocols 15, 2247–2276 (2020).

30. de Villarreal, J.M. et al. Pseudoalignment tools as an ejicient alternative to detect repeated transposable elements in scRNAseq data. Bioinformatics 39, btac737 (2023).

31. Feschotte, C. Transposable elements and the evolution of regulatory networks. Nature Reviews Genetics 9, 397–405 (2008).

32. Sundaram, V. & Wysocka, J. Transposable elements as a potent source of diverse cis-regulatory sequences in mammalian genomes. Philosophical Transactions of the Royal Society B 375, 20190347 (2020).

33. Nishihara, H. Transposable elements as genetic accelerators of evolution: contribution to genome size, gene regulatory network rewiring and morphological innovation. Genes & Genetic Systems 94, 269–281 (2019).

34. Rebollo, R., Romanish, M.T. & Mager, D.L. Transposable elements: an abundant and natural source of regulatory sequences for host genes. Annual review of genetics 46, 21–42 (2012).

35. Ohtani, H. & Iwasaki, Y.W. Rewiring of chromatin state and gene expression by transposable elements. Development, Growth & DiHerentiation 63, 262–273 (2021).

36. Zhang, Y., et al. Evolutionary rewiring of the wheat transcriptional regulatory network by lineage-specific transposable elements. Genome Research 31, 2276–2289 (2021).

37. Colonna Romano, N. & Fanti, L. Transposable elements: major players in shaping genomic and evolutionary patterns. Cells 11, 1048 (2022).

38. Borowsky, A.T. & Bailey-Serres, J. Rewiring gene circuitry for plant improvement. Nature Genetics 56, 1574–1582 (2024).

39. Craggs, J., et al. Inducing broadcast coral spawning ex situ: Closed system mesocosm design and husbandry protocol. Ecology and evolution 7, 11066–11078 (2017).

40. Cheng, H., Concepcion, G.T., Feng, X., Zhang, H. & Li, H. Haplotype-resolved de novo assembly using phased assembly graphs with hifiasm. Nature methods 18, 170–175 (2021).

41. Seppey, M., Manni, M. & Zdobnov, E.M. in Gene prediction: methods and protocols 227–245 (Springer, 2019).

42. Brůna, T., Hoj, K.J., Lomsadze, A., Stanke, M. & Borodovsky, M. BRAKER2: automatic eukaryotic genome annotation with GeneMark-EP+ and AUGUSTUS supported by a protein database. NAR genomics and bioinformatics 3, lqaa108 (2021).

43. Wang, X., et al. Draft genomes of the corallimorpharians Amplexidiscus fenestrafer and Discosoma sp. Molecular Ecology Resources 17, e187–e195 (2017).

44. Voolstra, C.R., et al. Comparative analysis of the genomes of Stylophora pistillata and Acropora digitifera provides evidence for extensive dijerences between species of corals. Scientific reports 7, 17583 (2017).

45. Zheng, G.X., et al. Massively parallel digital transcriptional profiling of single cells. Nature communications 8, 14049 (2017).

46. Hao, Y., et al. Integrated analysis of multimodal single-cell data. Cell 184, 3573–3587. e3529 (2021).

47 Alexa, A. & Rahnenführer, J. Gene set enrichment analysis with topGO. Bioconductor Improv 27, 1–26 (2009).

48. Sebé-Pedrós, A., et al. Cnidarian cell type diversity and regulation revealed by whole-organism single-cell RNA-Seq. Cell 173, 1520–1534. e1520 (2018).

49. Emms, D.M. & Kelly, S. OrthoFinder: phylogenetic orthology inference for comparative genomics. Genome biology 20, 1–14 (2019).

50. Stormo, G.D. Modeling the specificity of protein-DNA interactions. Quantitative biology 1, 115–130 (2013).

51. Liao, X., et al. A sensitive repeat identification framework based on short and long reads. Nucleic Acids Research 49, e100-e100 (2021).

52. Storer, J., Hubley, R., Rosen, J., Wheeler, T.J. & Smit, A.F. The Dfam community resource of transposable element families, sequence models, and genome annotations. Mobile DNA 12, 1–14 (2021).

53. Melsted, P., et al. Modular, ejicient and constant-memory single-cell RNA-seq preprocessing. Nature biotechnology 39, 813–818 (2021).

54. Quinlan, A.R. & Hall, I.M. BEDTools: a flexible suite of utilities for comparing genomic features. Bioinformatics 26, 841–842 (2010).

