## Supplementary_Files for "Cell-type-specific transposable element transcription tracks symbiosis and calcification programs in the reef-building coral *Acropora hemprichii*": Supplementary_figure.pdf

### Supplementary Figures

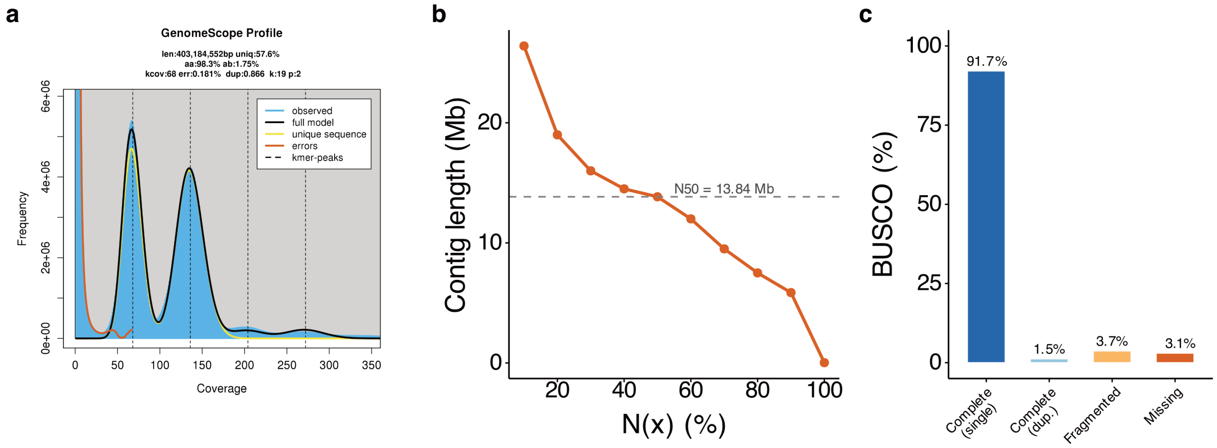


Fig. S1: BUSCO completeness assessment of the genome assembly of *Acropora hemprichii*. a. Genome size estimation based on k-mer analysis. **b**, Assembly contiguity assessed by the Nx curve. **c**, Genome completeness assessed by BUSCO analysis.

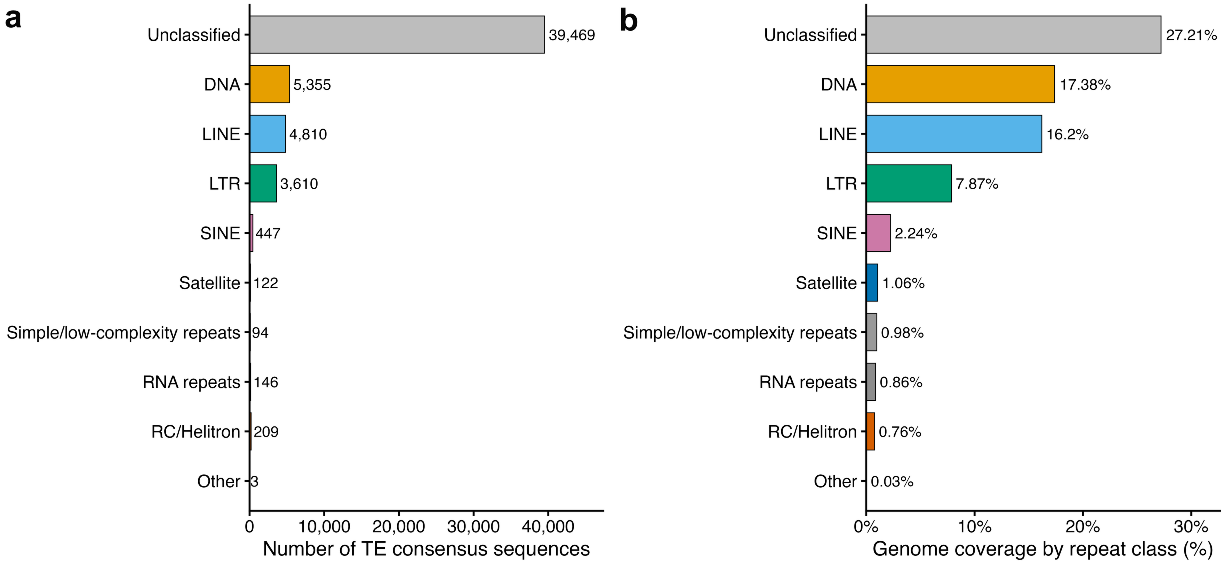


Fig. S2: Repeat annotation and genomic distribution of repetitive elements. **a,** Number of RepeatModeler-derived TE consensus sequences grouped by repeat class. **b,** Genome proportion annotated by RepeatMasker for each repeat class.

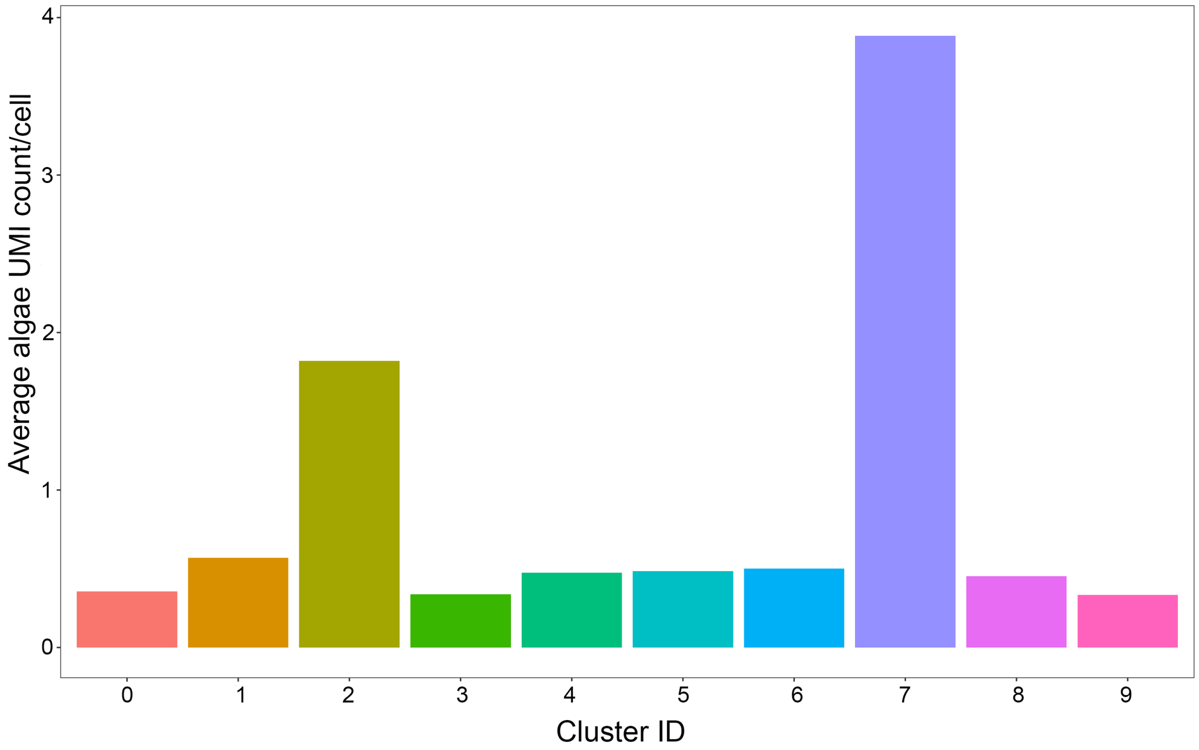


Fig. S3: Average UMI count from algae per cell within each identified cluster.

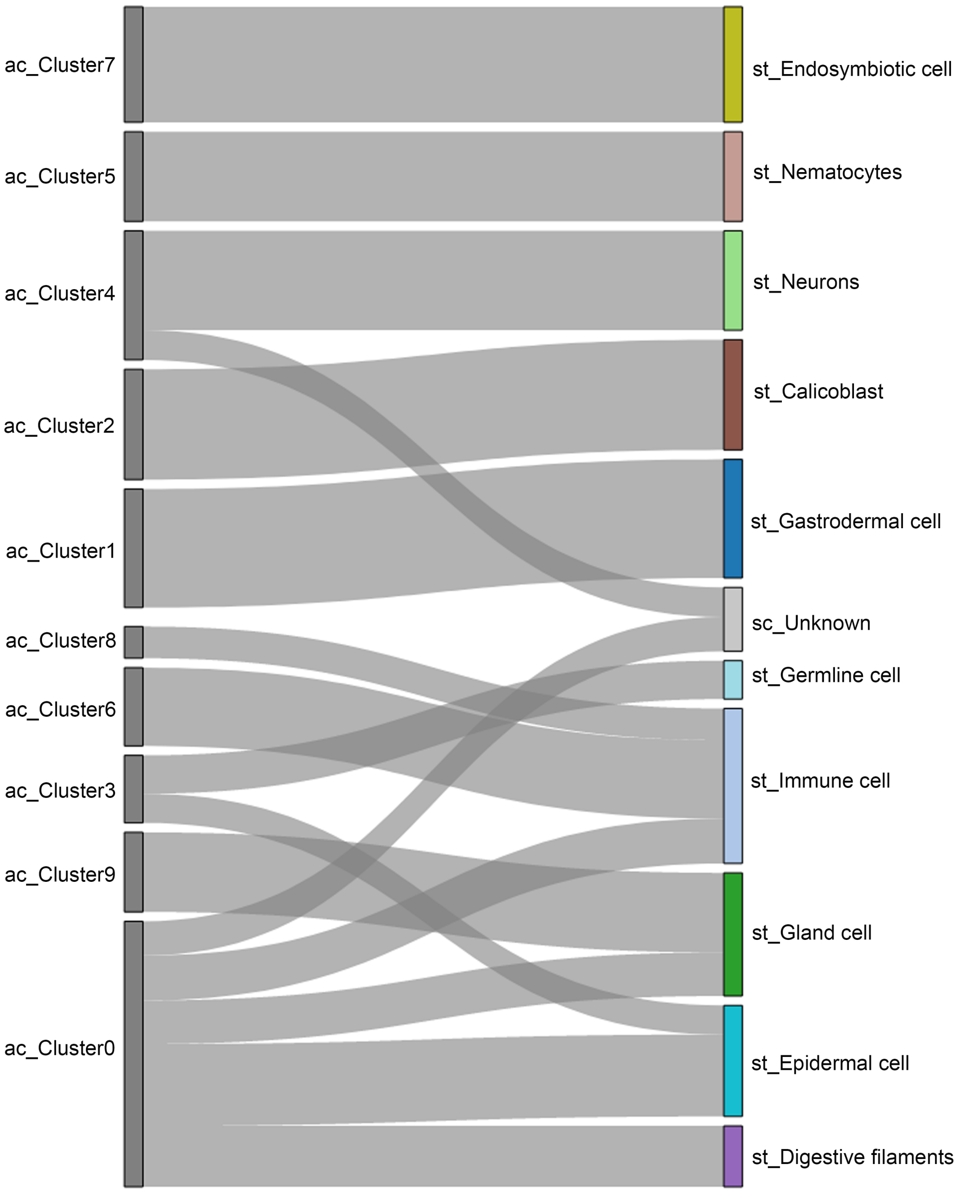


Fig. S4: Sankey plot summarizing the cell type mappings between *Stylophora pistillata* (st) and *Acropora hemprichii* (ac) clusters. Edges with alignment score <0.2 are omitted.

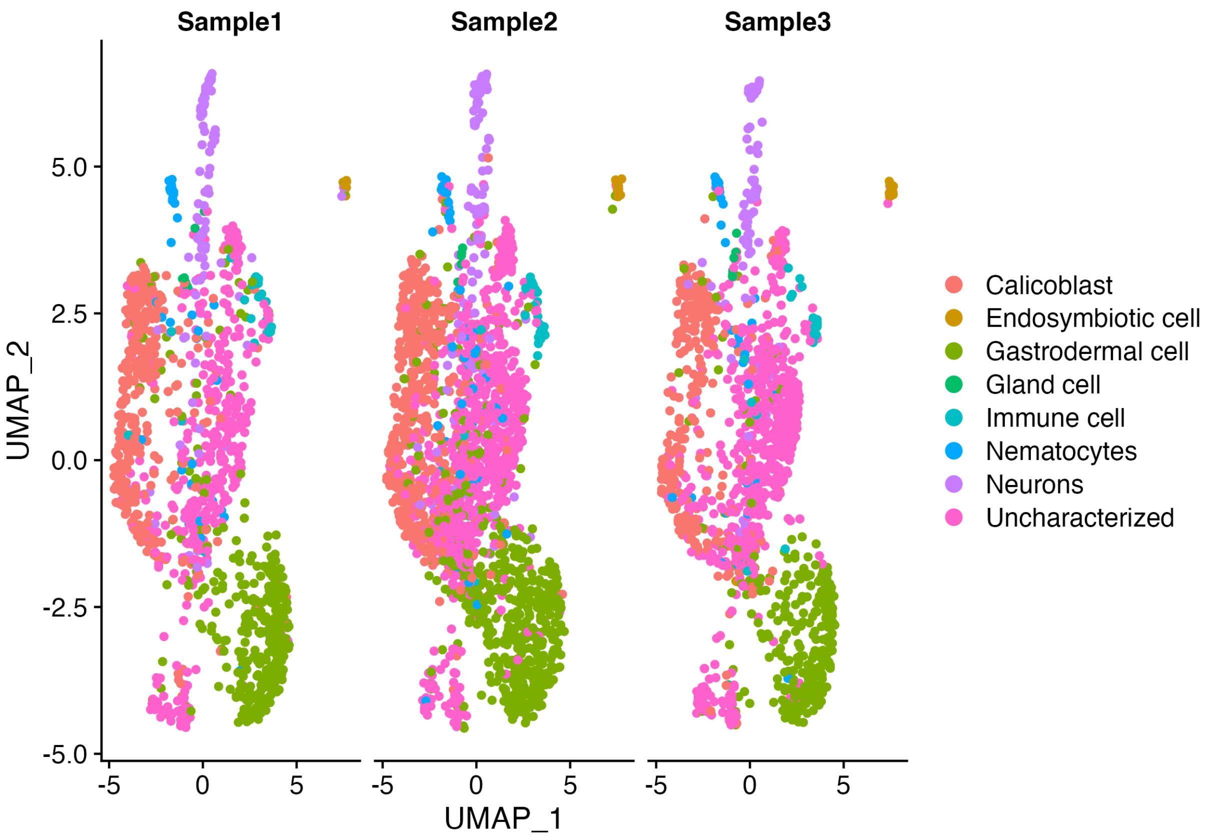


Fig. S5: UMAP plot of the transposable element expression by samples.

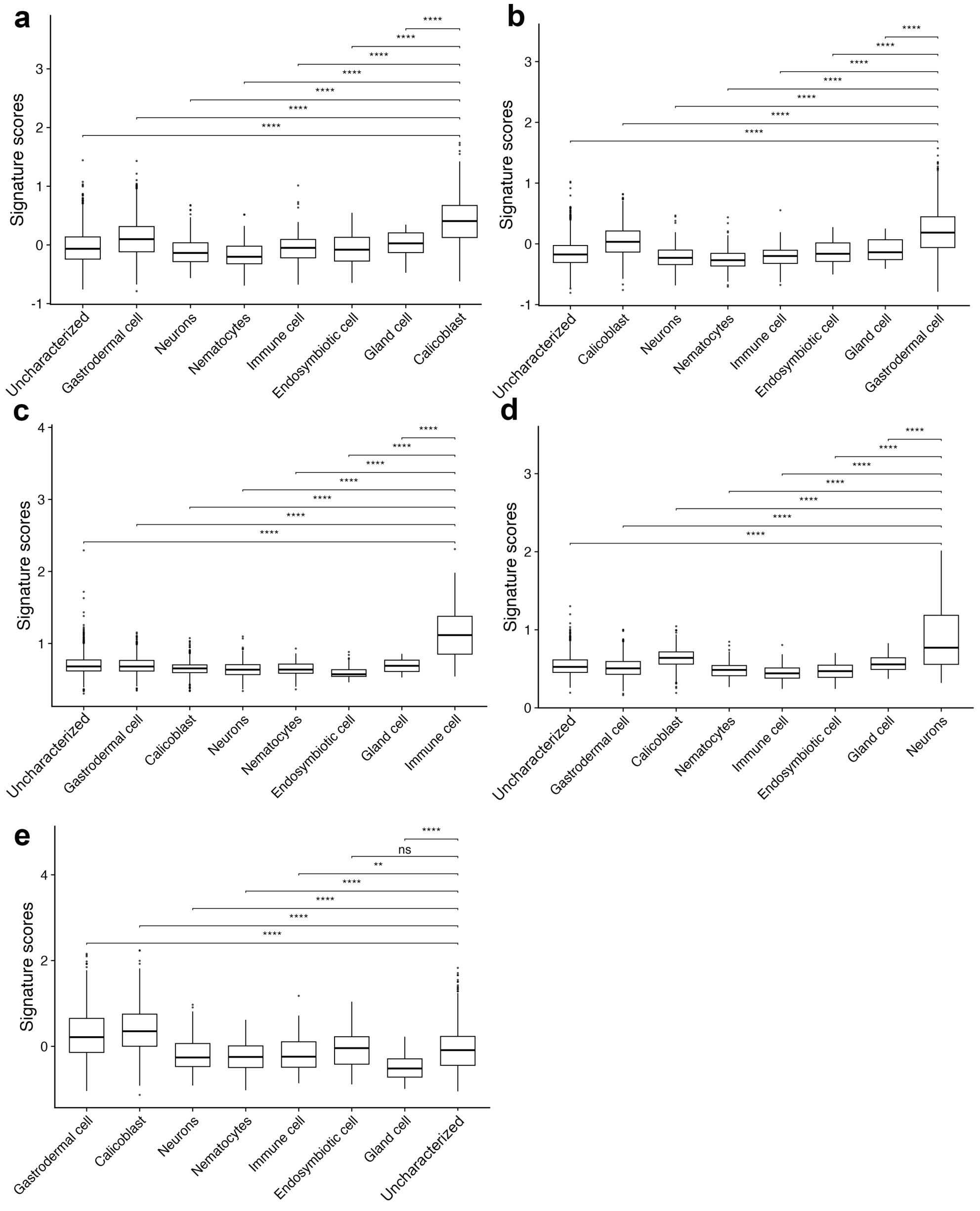


Fig. S6: Signature scores of the identified differential expressed TEs for calicoblast (**a**), gastrodermal cell (**b**), immune cell (**c**), neurons (**d**) and uncharacterized cells (**e**). A pair-wise t-test comparison is conducted between the target cell types and the others. ***: p-value < 0.001, **: p-value < 0.01, *: p-value < 0.05.

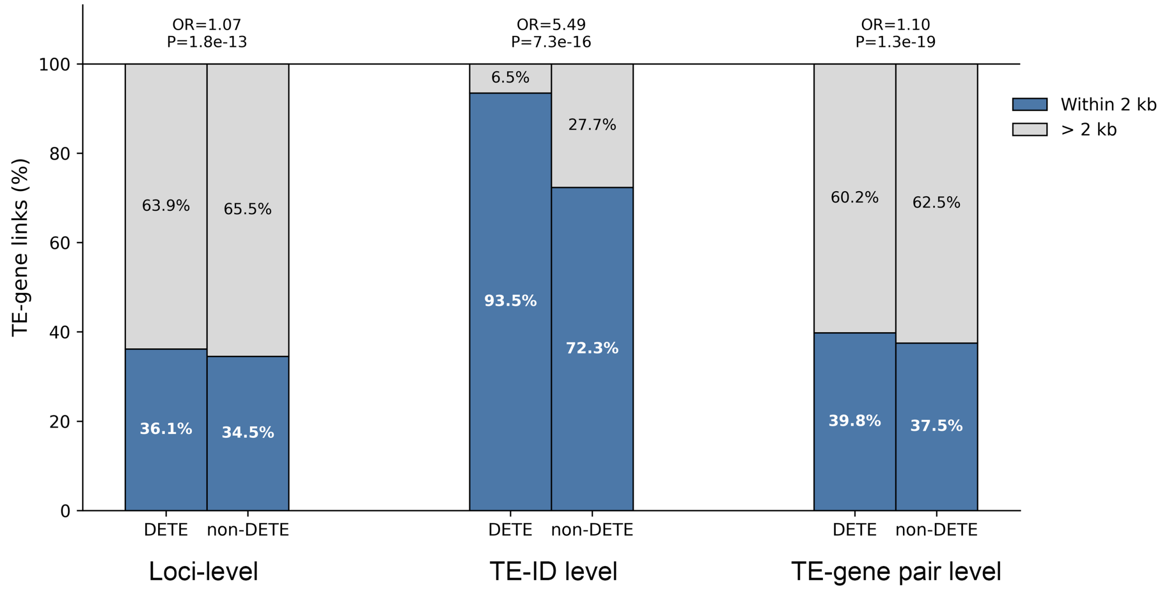


Fig. S7: TE–gene proximity enrichment across analytical levels. Stacked bar plots showing the proportions of DETE- and non-DETE-associated TE–gene links located within or beyond 2 kb of the nearest gene at the locus, TE ID, and TE ID–gene ID pair levels. Odds ratios and p-values were calculated using Fisher's exact test comparing DETEs and non-DETEs within each level.

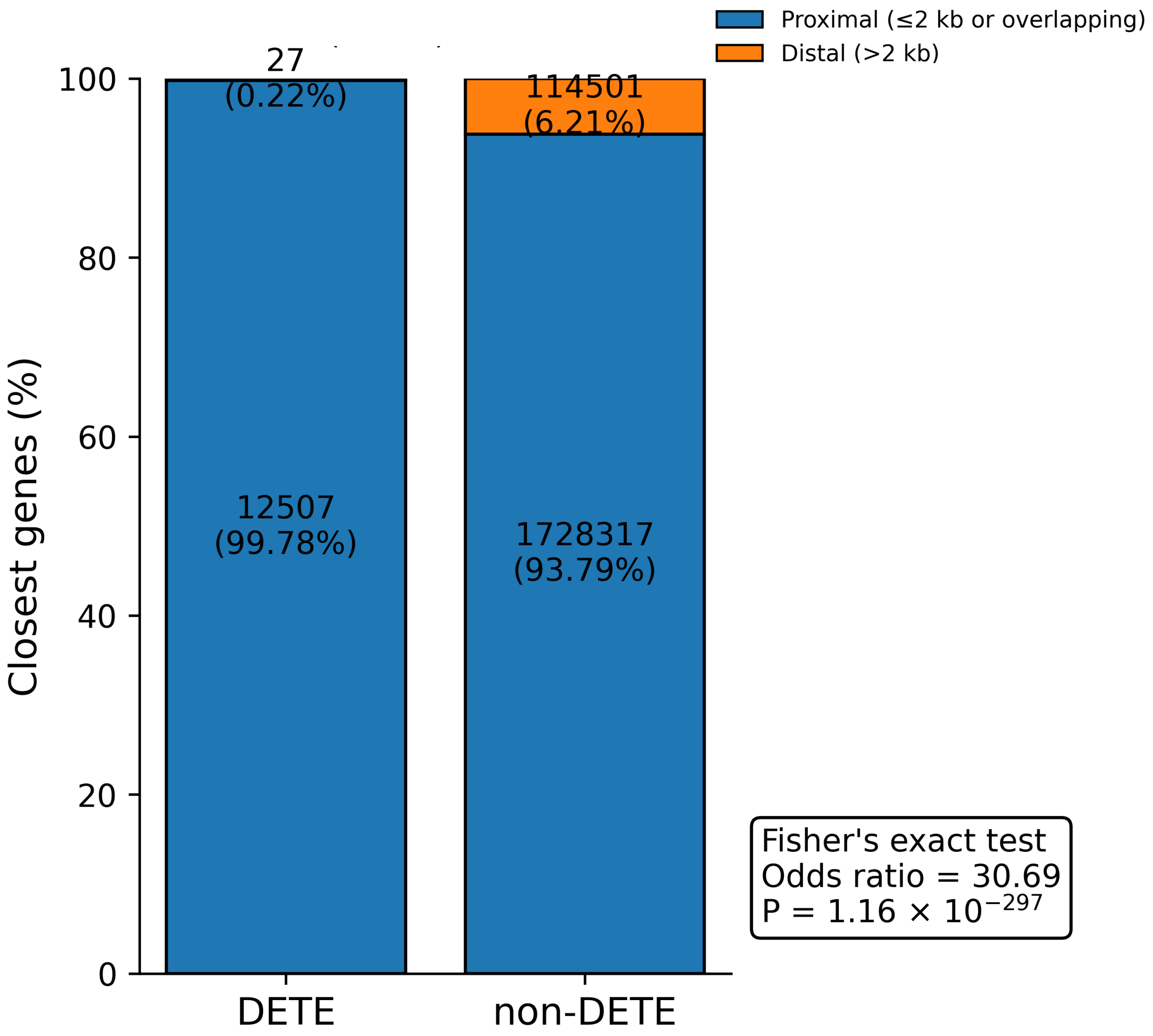


Fig. S8: Proximity enrichment of cell type–specific DETE-associated closest-gene assignments. Stacked bar plots showing the proportion of DETE- and non-DETE-associated cluster–TE closest-gene assignments classified as proximal (≤2 kb or overlapping) or distal (>2 kb). Odds ratio and p-value were calculated using Fisher's exact test comparing DETEs with non-DETEs.
